# A robust and efficient DNA storage architecture based on modulation encoding and decoding

**DOI:** 10.1101/2022.05.25.490755

**Authors:** Xiangzhen Zan, Ranze Xie, Xiangyu Yao, Peng Xu, Wenbin Liu

## Abstract

Thanks to its high density and long durability, synthetic DNA has been widely considered as a promising solution to the data explosion problem. However, due to the large amount of random base insertion-deletion-substitution (IDSs) errors from sequencing, reliable data recovery remains a critical challenge, which hinders its large-scale application. Here, we propose a modulation-based DNA storage architecture. Experiments on simulation and real datasets demonstrate that it has two distinct advantages. First, modulation encoding provides a simple way to ensure the encoded DNA sequences comply with biological sequence constraints (i.e., GC balanced and no homopolymers); Second, modulation decoding is highly efficient and extremely robust for the detection of insertions and deletions, which can correct up to ~40% errors. These two advantages pave the way for future high-throughput and low-cost techniques, and will kickstart the actualization of a viable, large-scale system for DNA data storage.

## Introduction

Synthetic DNA has now been proved to be a new potential storage medium for the exponentially growing data ^1, 2^. The total amount of data stored in synthetic DNA has reached the GB level; various practical automated read/write technologies for DNA storage have been proposed^3–5^. Unlike traditional electric/optical/magnetic storage media, DNA storage is characterized by large amount of insertions, deletions, and substitutions(IDSs) due to high error prone DNA synthesis and sequencing processes^6^. The difficulty of data recovery mainly comes from the synchronization problem because of random insertions and deletions. It is estimated that the error rate of the third-generation synthesis and sequencing technologies is about 10–15%^7^. For example, Antkowiak et.al^8^ found that in the 15 million retrieved sequences, over 99.9% were retrieved with error. Lee Organick^9^ reported that over 88% of reads were of incorrect length because of the insertions and deletions caused by synthesis and sequencing. Additionally, sequence loss can also occur during synthesis, amplification, and sequencing. The complex IDS errors and sequence loss present a daunting challenge to the reliable recovery of stored data.

To reduce the occurrence of various errors, most of the prior works complied with some encoding constraints on DNA sequences, such as no homopolymers, guanine–cytosine (GC) content of 40% ~ 55%, and no secondary structures. For example, Goldman et al. ^10^adopted a 3-base rotation encoding scheme. Randomization with a pseudorandom binary sequence is a popular strategy utilized to avoid homopolymers by many works^9, 11^. However, there is not yet a universally accepted standard to build upon for large scale storage applications.

There is a variety of literatures dedicated to the study of reliable data recovery from the distorted sequenced reads by the IDS channel of DNA storage. Davey and MacKay^12^ were the first to propose a probabilistic model for the binary IDS channel where the maximum a-posteriori (MAP) reconstruction was inferred by a hidden Markov model (HMM). To take advantage of the multiple copies of each original strand, Lenz et. al. proposed two decoding algorithms for multiple received sequences based on the HMM^13^. Later, Srinivasavaradhan et al. proposed a heuristic algorithm, Trellis BMA, which allows multiple single-trace trellis decoders to interact and estimate soft output^6^. Different from these HMM-based method, Yadiz et al.^14^ developed an integrated pipeline which gradually corrects the IDSs in multiple phases. It consists of a multiple sequence alignment (MSA) module to obtain the consensus sequence, an iterative Burrows-Wheeler alignment (BWA) module to improve the reconstruction quality, and a post processing module to refine the sequence by some encoding constraints. Antkowiak et. al^8^ applied MSA to solve most of the IDSs, and the remaining substitutions in the consensus sequence were further corrected by the inner decoder. The decoding issue can also be considered as a searching problem. Press et. al.^15^ proposed a special convolution code, HEDGES, which could use A* algorithm to search for the possible path, and other failure path were regarded as erasures to be corrected in the outer code. Song et al.^16^ constructed a de Bruijn graph (DBG) using multiple sequences and infer the correct sequence by a graph searching algorithm. For English text storage, our group constructed a 6-base code table for characters and the IDSs were corrected by distance-based code correction, multiple sequence voting, and spelling check^17, 18^. Additionally, Xue et al.^19^ proposed a special Levenshtein code^20^ capable of correcting a single synchronization or substitution error in a code word. In short, the error correcting capacity of current methods ranges from 5-18% with various logic densities (≤2bit/nt).

Sequence loss may be somewhat alleviated through increasing sequencing coverage. However, it is not sufficient for lossless data recovery in large-scale storage. In the pioneer work of Church et.al^21^, sequence loss was observed even in sequencing coverage 3000×. DNA Fountain ^22, 8^ and inner-outer code were proved to be an effective way to overcome this issue.

In the future, large scale application of DNA storage claims synthetic biotechnologies with higher throughput but lower cost ^8^. This means we have to face an even more error prone storage situation. Meanwhile, stored data by DNA may suffer from some unpredicted or malicious damages, like the hard disk failures in computers. In addition, DNA molecules may undergo degradation and break over time. To pace with these challenges, it is necessary to develop more robust techniques which can tolerate higher error settings (e.g. 20~30%).

In the field of telecommunication, modulation has accomplished reliable signal transmission by superimposing baseband signals (or low frequency signals) to the carrier signals (or high-frequency signals) ^23–25^. The modulated signal not only has the power to anti-interference, but also has no effect on the modulating signal. In fact, the carrier signals serve as the backbone to protect the modulating signal from external disturbances. In this paper, we develop a new DNA storage architecture based on modulation encoding and decoding. Results from computer simulation and real data demonstrate that this storage architecture has three advantages: First, modulation encoding provides a storage friendly encoding architecture to satisfy biological constraints and have similar thermodynamic property. Second, modulation provides an effective way to solve synchronizations, and the proposed method can tolerate up to ~40% errors. The decoding process is very time efficient and suitable for large scale application. To the best of our knowledge, this architecture far exceeds the comprehensive performance of any state-of-the-art works.

## Results

### Modulation-based DNA storage

We assume that clustering is already completed and we focus on the reconstruction problem of the clustered sequences. Fig.1a shows the four steps of the proposed modulation-based storage paradigm. First, the original binary information is modulated with the carrier stream to generate DNA sequences, according to a predefined rule. Second, the observed strands are transformed into marked reads by aligning their observed carriers with the carrier strand to solve the synchronizations. Third, align the marked reads in each cluster and use majority voting to obtain a consensus DNA sequence. Finally, use the carrier strand to demodulate the consensus DNA sequences to recover the original binary information.

**Fig. 1.**
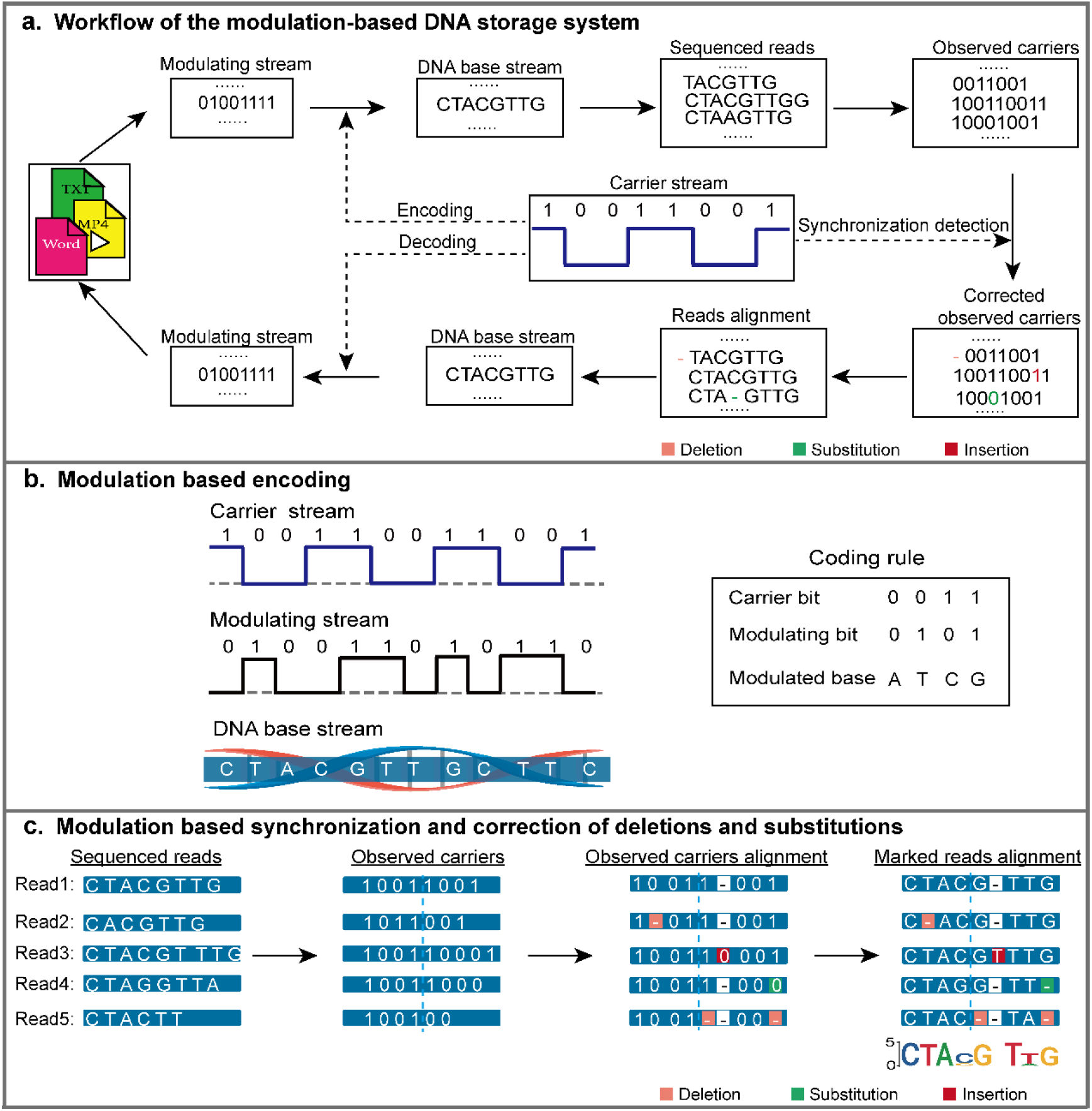
Overview of modulation-based DNA storage. **a**, Workflow of encoding and decoding processes. **b**, Modulation-based encoding. **c**, Modulation-based synchronization and correction of deletions and substitutions.

### Encoding with the carrier strand

In DNA storage, the stored binary information is usually partitioned into many strands with a fixed length *n*. All these strands are modulated with a common carrier strand *c* to generate their corresponding DNA sequences. In this paper, the carrier strand *c* is composed of repeated substring like ‘1001’, whose length (carrier period) is denoted as *p_c_*. Given a binary carrier strand *c* =‘**1001**1001**1001**’, a binary message strand *m* =‘010011010110’ is transformed bit by bit into a DNA sequence *s* =‘**CTAC**GTAG**CTTC**’, according to the following rule (shown in Fig. 1b):

1. If *c*[*i*]=‘0’ (0 < *i* < *n*), it will modulate *m*[*i*]=‘0’ to ‘A’ and *m*[*i*]=‘1’to ‘T’;
2. Otherwise*c*[*i*] =‘ 1 ’, it will modulate *m*[*i*] =‘0’ to ‘C’ and *m*[*i*] =‘ 1 ’ to ‘G’.

Modulation encoding provides a convenient mechanism to satisfy the sequence constraints by selecting appropriate period of *c*. Not only can we easily strike a balance in GC content, but it can also be uniformly distributed across the sequence. For example, given *c* =‘**1001**1001**1001**’, all encoded sequences will share a similar pattern of two A/T bases surrounded by a pair of G/C bases. DNA sequences with this pattern have desirable biological properties, such as similar melting temperatures and homopolymers of at most 2.

### Decoding with the carrier strand

The main challenge in decoding comes from the synchronization issue, where insertions and deletions (indels) cause bases to deviate from their original positions. Given *t* reads *r*_1_,…,*r_t_* for a sequence *s*, the decoding problem can be formulized as inferring a consensus sequence *s*\* with the maximal posteriori likelihood *p*(*s*\*|*r*_1_,…,*r_t_*). We divide this problem into two subproblems: synchronization and correction of deletions and substitutions.

We first solve synchronization based on the carrier strand. According to the modulation rule, any read *r_k_* of sequence *s* can be transformed to a binary strand 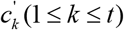, which we call the observed carrier strand. Obviously, 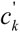 should be similar to *c* in a large degree as *s* is produced by the carrier strand *c*. Therefore, the optimal alignment of 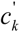 to *c* should discover the occurrences of IDSs in read *r_k_*. Taking *c* =‘10011001’ and *r_k_* =‘CTAATAG’ as an example, the best alignment of 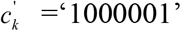 to *c* should be ‘100**0**-001’ which includes one substitution in the 4^th^ position and one deletion in the 5^th^ position (‘-’ denotes deletion). This means that *r_k_* may involve one substitution and one deletion in the corresponding positions. That is, it should be modified as 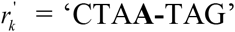, which we name as the marked read of read *r_k_*.

We then correct the deletions and substitutions in the marked reads 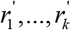 by majority voting. After synchronization, the marked reads are all have the same length with the original sequence *s*, and only deletions and substitutions are existed in them. These errors can be simple solved by a majority voting in each position *i*. Fig. 1c shows the main steps of the modulation-based error detection and correction (orange for deletion, red for insertion and green for substitution).

Finally, the original binary message can be simply obtained by demodulating the obtained consensus sequence using the opposite of the encoding rule. If *c*[*i*] =‘0’, ‘A’ will be demodulated to *m*[*i*] =‘0’ and ‘T’ to *m*[*i*]=‘1’; otherwise, ‘C’ will be demodulated to *m*[*i*] =‘0’ and ‘G’ to *m*[*i*] =‘ 1 ‘.

In sum, the carrier signal *c* provides valuable prior knowledge for all encoded sequences. This information proves to be highly useful in solving the synchronization problem based on the pairwise alignment of a distorted 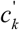 to the template *c*. As we know, sequence alignment has been well studied in bioinformatics, and many practical tools, such as MAFFT^26^, Clustal Omega^27^, Coffee^28^, and MUSCLE^29^, have been developed based on the famous Needleman-Wunsch algorithm^30^. In this paper, we apply MAFFT, an efficient MSA software, to 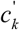 and *c*. Its time complexity of pairwise alignment is reduced from O(*n*^2^) to O(*n*log*n*) by the Fast Fourier transform (FFT). Therefore, the read synchronization process can be accomplished in quasi-linear time O(*n*log*n*).

### Sequence properties of modulation encoding

To measure our encoding strategy against the classical methods, we compare the distributions of GC content, maximum homopolymers, and melting temperatures of the encoded DNA sequences. These four classical methods are Base Coding(00->A, 01->C, 10->G, 11->T), Goldman 2013^10^, Church 2012^21^, and DNA Fountain 2017^22^ (shown in Fig. 2). For our method, the carrier strand *c* =‘1001……1001’ is used to encode the original information.

**Fig. 2.**
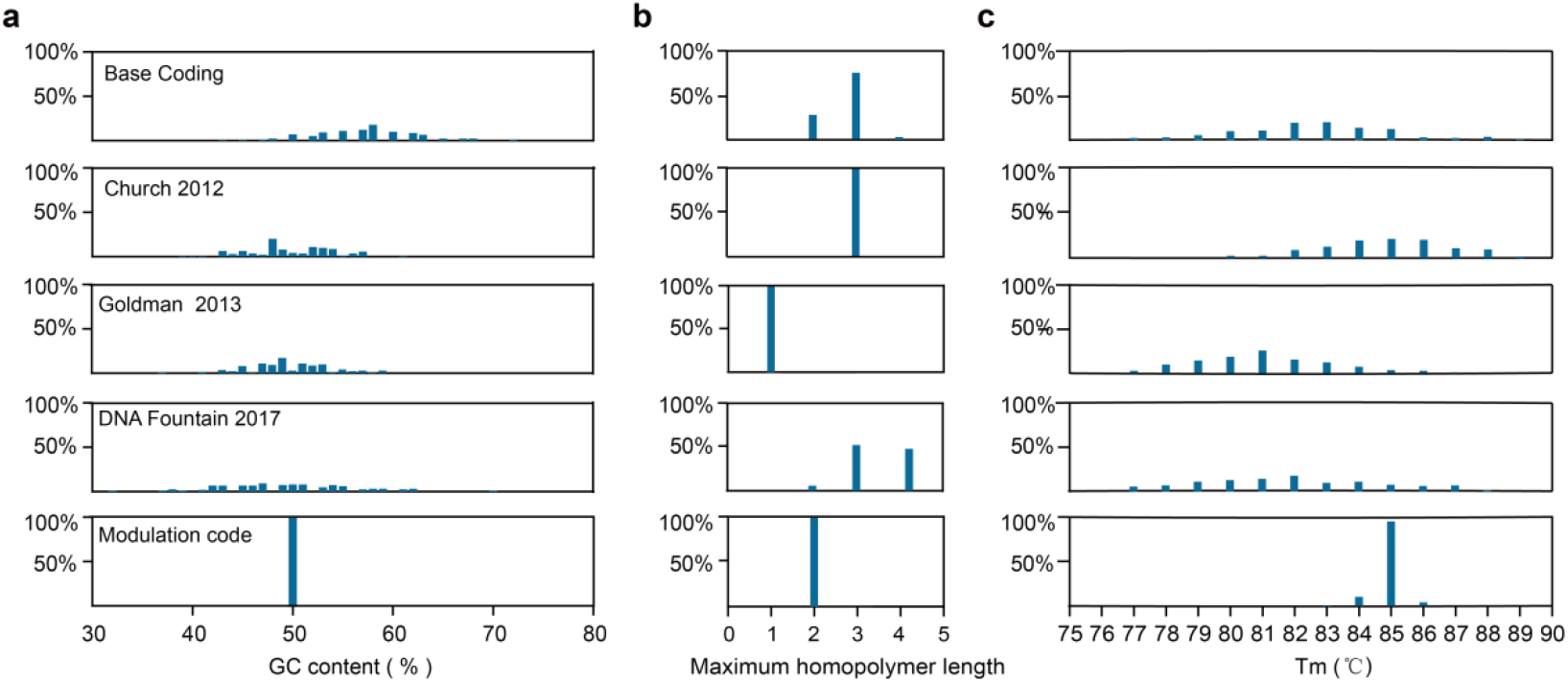
Comparison of the properties of DNA sequences by Base Coding, Goldman 2013, Church 2012, DNA Fountain code 2017 and modulation-based encoding. **a**, Distribution of GC content. **b**, Distribution of maximum homopolymers. **c**, Distribution of melting temperatures Tm.

We can see that sequences by modulation have fixed GC content while others generally range between 30~70% (Fig. 2a). The maximal homopolymers in sequences by Goldman 2013^10^ and modulation are 1 and 2 while other may even reach 4 (Fig. 2b). These two observations demonstrate that by selecting an appropriate carrier strand *c*, modulation encoding provides a convenient mechanism to satisfy the sequence constraints without any preprocessing adopt by previous methods^9, 11^.

To predict the melting temperature of sequences, we apply a web tool MFEprimer (https://mfeprimer3-0.igenetech.com/) ^31^. The melting temperature of sequences encoded by modulation, is around 85°C while others range in 77~87°C (Fig. 2c). The stable thermodynamic property is mainly attributed to the uniform distribution of GC across the sequences. This demonstrates modulation encoding can accomplish a storage friendly encoding mode which is favorable to DNA synthesizing, PCR (Polymerase Chain Reaction), and sequencing processes. Not only can this feature reduce the occurrences of unexpected errors to some extent, but it can also improve the efficiency of the data reading processes (i.e., PCR and sequencing).

### Decoding performance using different carrier periods

Intuitively, longer period carrier strands tend to include more complex patterns, and such complex patterns may help to detect abnormal indels. One way to describe the complexity of sequences is to count the number of substrings in them. For example, carrier strand *c*_1_= ‘10101010’ include substrings ‘10’, ‘01’, ‘101’, ‘010’, ‘1010’, and ‘0101’, while *c*_2_= ‘10011001’ includes substrings ‘10’, ‘00’, ‘01’, ‘11’, ‘100’, ‘001’, ‘011’, ‘110’, ‘1001’, ‘0011’, ‘0110’, and ‘1100’. Obviously, the later contains more substrings than the former. Fig. 3 shows the average decoding performance by carrier strands with period 2, 4, 8, and 16 respectively.

**Fig. 3.**
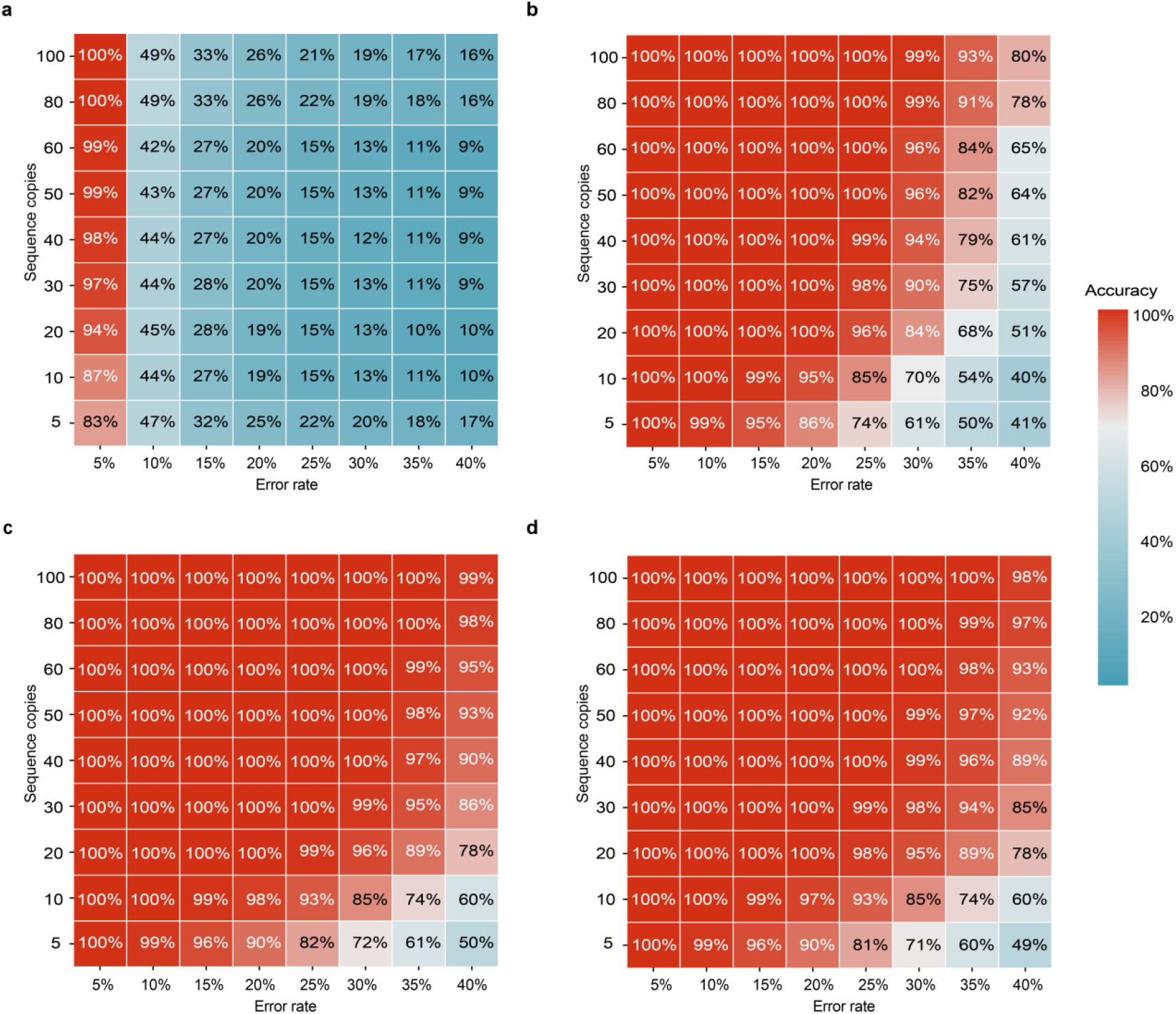
Decoding performance at different carrier periods. **a**, *p_c_* = 2. **b**, *p_c_* = 4. **c**, *p_c_* = 8. **d**, *p_c_* = 16. The value in each colored box denotes the average recovering accuracy at the corresponding error rate and sequence copies.

First, the average decoding performance tends to be stable as the period increases. When the period is 2, complete recovery can only be accomplished in the lowest error rate we tested, and the average performance is below 50% in other error settings. However, the average performance improved dramatically as the period increases to 4. And it becomes stable as period increases to 8. These observations concur our assumption that longer period is beneficial to complicate error settings. However, longer periods may affect the homogeneous distribution of GC content and the control of homopolymers. Therefore, considering both the sequencing property and the error correction performance, we find that setting the period to 8 may be appropriate in most situations.

Second, modulation-based decoding with period 8 and 16 is extremely robust to IDSs. At low error rate levels, such as 5%, it can completely recover the data with just 5 sequence copies, which is far less than the minimal copies required in previous works. Even when the error rate is as high as 40%, it can recover about 99% of the data given 100 sequence copies. However, the required sequence copies for complete recovery increases dramatically when the error rate is higher than 30%. One major reason is that the combination of deletions and substitutions at certain positions requires additional sequences for correction. To the best of our knowledge, this performance far exceeds the error correction capability of any state-of-the-art methods with a logical density 1bit/nt (See Table 1). Such performance mainly stems from its powerful ability to solve the synchronization problem using the carrier strand. As the decoding process includes two individual steps: synchronization and correction of deletions and substitutions, the error correction process can sufficiently take advantage of the information in the multiple reads. This strategy is different from the previous methods, which apply HMM^6, 13^, MSA^8, 14^, GPS^16^, or A*^15^ algorithms to solve all three errors at the same time. Our result demonstrates that not only can solving the synchronization problem first simplify the complexity of the decoding process, but it can also significantly improve error tolerance.

**Table 1.**
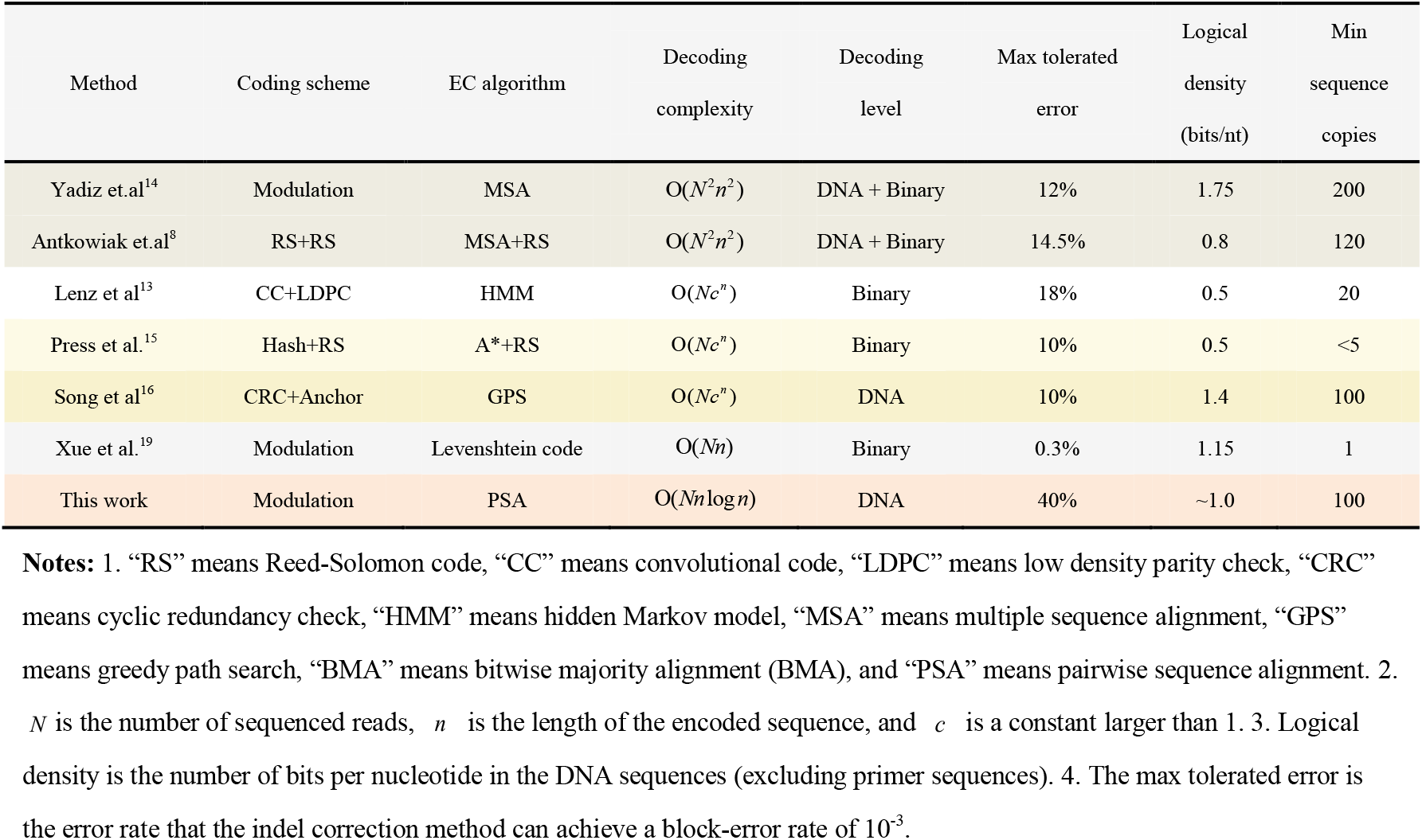
Comparison with the state-of -the-art methods.

### Decoding performance for consecutive insertions/deletions

Previous studies have demonstrated that consecutive indels are frequently observed in sequenced reads, which may result in many abnormal reads with incorrect lengths ^7, 32, 33^. Compared with a single insertion or deletion, consecutive indels are more difficult to correct. For example, HEDGES can deal with consecutive deletions, but can’t tolerate more than 2 consecutive insertions^15^. We further investigate the decoding performance of the proposed method on consecutive indels. For simplicity, we assume that insertion or deletion errors in all reads occur consecutively and have the same length.

Fig. 4 shows the average decoding performance with period 8 and 16 given sequence copies 30 and 60, respectively. As the consecutive length increases, the performance on period being 16 is better than that of period 8, especially for error rate ≥ 20%. As the error rate increases, longer consecutive indels may destroy the periodic structure of the carrier strands, which may affect its error detection capability. This is the main reason that carrier strands with period 16 are more robust than those with period 8, as the lengths of consecutive indels are far less than 16. In addition, increasing the sequence copies can improve the decoding performance at high error rates.

**Fig. 4.**
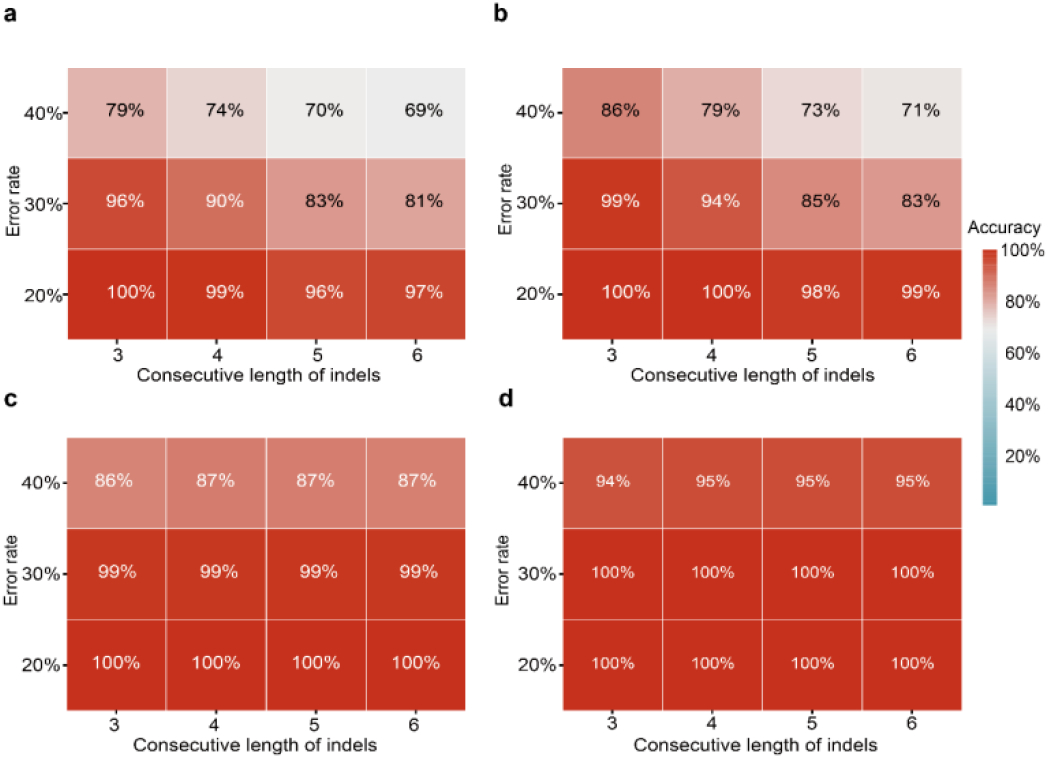
Decoding performance to consecutive insertions/deletions. **a**, period 8, sequence depth 30. **b**, period 8, sequence depth 60**. c**, period 16, sequence depth 30. **d**, period 16, sequence depth 60.

In most previous works, low-quality reads are usually discarded, which for large-scale application may lead to non-negligible loss both in cost and time. However, this problem can be significantly alleviated as modulation-based error detection can defend against the consecutive indels.

### Decoding Performance on a Real Dataset

We further illustrate the performance of the proposed method on a real dataset published by the Microsoft group. The original information was first added some marker repeat (MR) codes and then translated into DNA sequences according to the quaternary code (00-A, 01-T 10-G, 11-C). To apply the proposed method, we first construct a carrier strand for each DNA sequence by translating A/T to 0 and G/C to 1. Then the decoding process is used for the reads in each cluster.

Fig. 5a shows the average normalized Hamming distance between the decoded information and the original one under different sequence copies. Clearly, the proposed method (red color) dramatically outperforms the Trellis BMA^6^(blue color) and the multiply posteriors method (orange color) by Lenz et al.^13^. This indicates our modulation-based decoding method is superior to the other two HMM-based methods.

**Fig. 5.**
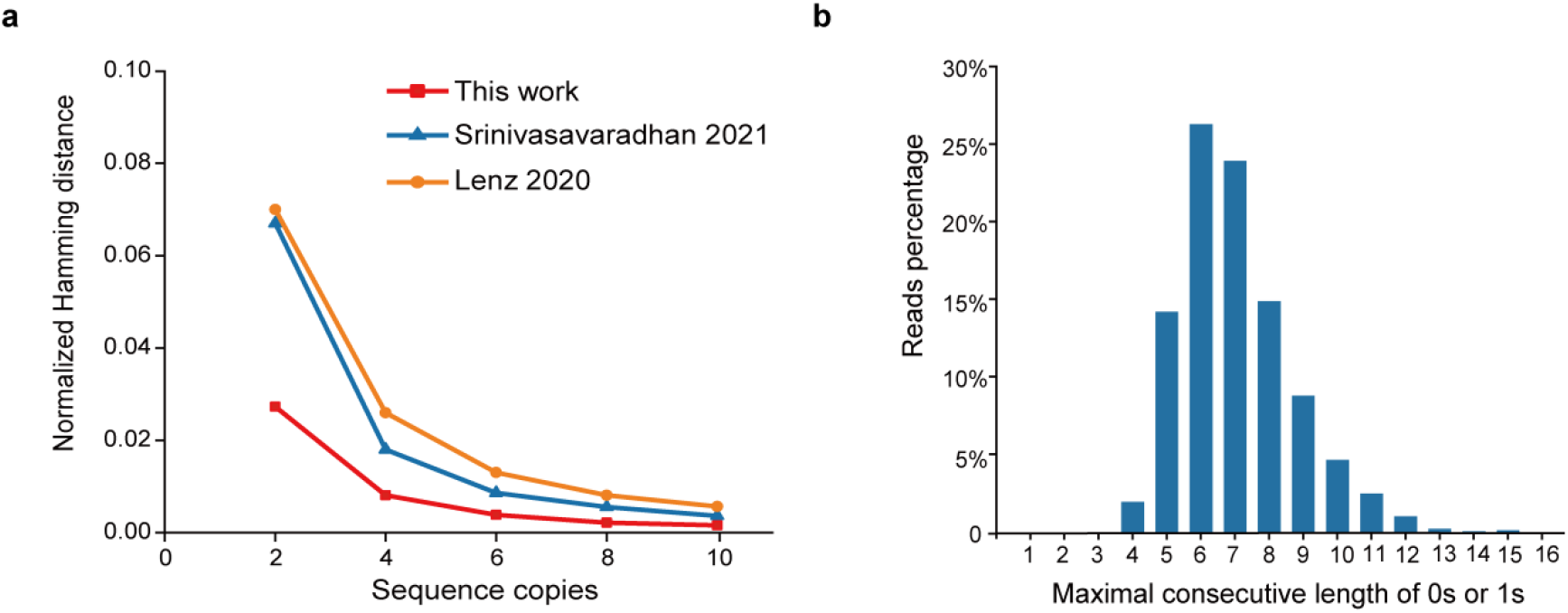
The performance of our proposed method on real data with the method of Lenz et al. ^13^ and Srinivasavaradhan^6^ et al. **a**, The normalized Hamming distance at sequence copy 2,4, 6, 8, 10. **b**, Distribution of the maximal consecutive 0s or 1s in the carrier strands of the encoded DNA sequences.

Although the estimated error rate of the dataset is about 5.9%, there still exist a few uncorrected errors with sequence copies being 10 by our method. To investigate this inconsistency with the results in Fig. 3, we further check the maximal consecutive 0s or 1s in the constructed carrier strands. Fig. 5b. shows that more than 80% of the carrier strands have at least 6 consecutive 0s or 1s. That is, these encoded DNA sequences include many continuous GC or AT regions. Such consecutive 0s or 1s may reduce the error detection capability of the carrier strands. This further verifies that simple patterns may limit the carrier strands’ error detection capability.

## Discussions

To take a comprehensive review, Table 1 lists the coding scheme, error correction (EC) algorithm, time complexity, reported maximal tolerated error rate, and logical density of six classic works and ours. We give a brief discussion in the following aspects.

Our method’s encoding is simple and storage friendly. By selecting the appropriate carrier strand, the encoded DNA sequences not only satisfy the constraints of GC content and homopolymers, but also have similar thermodynamic properties, which are beneficial to the biochemical techniques and may help to reduce errors. However, other encodings have to take hash or convolution operations^15^, add RS/LDPC/CRC redundancy^8, 13^, or perform XOR randomization on the original binary stream before translating them into DNA sequences. Fountain code^22^ and the de Brujin graph method by Song et.al ^16^ both need a filter process to discard binary streams containing illegal subsequences. Moreover, it should be noted that Yadiz et.al^14^ and Xue et al.^19^ also adopted a modulation encoding which were essentially different from ours. They used two information strands to produce DNA sequences according to the encode rules we used. Therefore, it has nothing to do with signal modulation and solving the synchronization issue as we do.

Our method can tolerate up to 40% errors, which far exceeds the state-of-the-art methods. The methods by Lenz et al. ^13^, Antkowiak et al. ^8^, and Press et al. ^15^ can only tolerate errors up to 18%, 14.5%, and 10%, respectively, with logical density <1. The Levenshtein code method by Xue et al.^19^ is only feasible to very low error situations (e.g. 0.3%). The methods by Song et al. ^16^ and Yadiz et al.^14^ have a higher logical density than ours: 1.5 and 1.75, but can only correct errors up to 10% and 12%. In coding theory, the more logical redundancy is added, the more powerful the error correction ability. To tolerate higher errors, these methods would have to add more logical redundancy, inevitably lowering their logical density. However, our method can achieve this by just increasing the sequence copies without sacrificing the logical density (See Fig. 3 C/D).

Our modulation-based synchronization is more powerful than the other previous methods. In our method, the alignment between the observed and the true carrier strand could optimally detect the occurrences of synchronization. In the MSA-based methods by Yadiz et.al^14^ and Antkowiak et.al^8^, synchronization is solved by the progressive alignments between similar reads. At low error regime, although the true conserved information is preserved in most of the common regions, there still exist some missed alignments caused by the random IDSs. Such situation may worsen as the channel error increases, which then limits the ability of solving synchronization at high error regime.

In the HMM-based method by Lenz et al.^13^ and Srinivasavaradhan et al.^6^, the watermark codes function similarly as the carrier strand in ours. However, their synchronization ability is limited by the channel noises and the transmitted information embedded in the sparse codes. In the work by Lenz et.al^13^, they could tolerate errors as high as 18%, but has a very low logical density of 0.5bit/nt. In addition, HMM model usually needs to accurately estimate the probability of IDSs and these parameters are dependent on the used techniques. This will hinder its application under different scenarios. In contrast, our method can be applied to any situation without considering the difference of the platforms.

In HEDGES by Press et. al. ^15^, synchronization is solved based on the associations between consecutive bits. The performance of A* algorithm will decrease dramatically as errors increases. The author claimed that their method could tolerate consecutive insertions at most 2.

The DBG-based method by Song et. al ^16^ attempted to find the correct path in a de Bruijn graph which was consisted of uncontaminated DNA k-mers. However, the probability that a k-mer is uncontaminated will drop dramatically as the error rate increases. For k=18 in their work, this probability drops from 15% to 5% as error rate increases from 10% to 15%. Although increasing sequence coverage may alleviate this effect to some degree, the probability of the correct path consisted of tens or hundreds of k-mers will tend to zero (0.05^303^, *n* =320 nt). Therefore, such graph searching based method may not work for error rate larger than 15%.

The decoding process of our method has is highly time-efficient and suitable for large scale application. Its most time-consuming process is synchronization, where pairwise sequence alignment (PSA) takes place. Therefore, its time complexity is O(*Nn* log *n*), where *n* and *N* are the number of the sequenced reads and their length, respectively. The method by Antkowiak et. al^8^ involves multiple sequence alignment (MSA) in each cluster and the normal RS decoding. It has a polynomial time complexity of order O(*Nn*^2^). The time complexities of the other three are determined by the A*, hidden Markov model (HMM), and greedy path search (GPS) algorithm, respectively. Although various heuristic strategies can be applied, they all have an exponential time complexity *O*(*Nc^n^*), where *c* > 1 is a constant determined by the average searching branches.

## Conclusion

At this point, the cost per bit using current DNA storage technologies is still much higher than those of traditional electronic and optical storage devices. Developing DNA storage-oriented technologies allowing more errors may provide enough room for further reducing the cost of synthesis and sequencing. Modulation-based DNA storage is characterized by storage-friendly encoding, ultra high error tolerance, and extreme efficiency in decoding. Therefore, it not only paves a solid foundation for reliable information retrieval in high error environment, but also could drive the development of low-cost synthetic technologies. We believe that this new storage architecture could facilitate the early realization of large-scale DNA storage application.

## Methods

### Datasets used for experiment results

For simulation experiments, a text file “The Grandmother” (excerpted from “Andersen’s Fairy Tales), is encoded into 140 DNA sequences of 120 bases (8 bases for index and 112 bases for data). Error rates in the encoded sequences range between 5% ~ 40%, where insertions, deletions, and substitutions are equally likely. The sequence coverage range from 5 ~ 100. All experimental results are obtained by repeating 1,000 times under a given error rate and a fixed number of sequence copies.

The real dataset^6^ includes 269,709 reads of 10,000 uniform random DNA sequences of length 110 (https://github.com/microsoft/clustered-nanopore-reads-dataset). All DNA sequences were synthesized by Twist Bioscience and sequenced using ONT MinION, and the estimated error rate in the sequenced reads is about 5.9% in total (p_ins_=1.7%, p_del_=2%, p_sub_=2.2%). The noisy reads were grouped by a pseudo clustering algorithm^34^.

### Construction of the periodic carrier strands

In this paper, we investigate the performance of carrier strands with period *p_c_* = 2, 4, 8, 16. To satisfy the constraints of GC content and maximal homopolymers, the period substrings in the carrier strands should satisfy the following two criterions:

1. The percentage of 1s (or 0s) should be 50%.
2. The consecutive length of 1s (or 0s) should be less than 4.

For period 2, there are only two carrier strands: ‘0101…0101’ and ‘1010… 1010’, which constitute of substrings ‘01’ and ‘10’. For period 4, there are 4 carrier strands: ‘0110’, ‘1001’, ‘1100’, and ‘0011’,. For period 8, we enumerate all binary strings with length 8, and discard those with period 2 and 4. Substrings for period 16 is obtained in the same way.

### Error-correction for sequenced reads

The proposed error-correction process is illustrated in Fig. 1c, and it contains the following steps: Step 1, for each reads cluster, derive the observed carrier strand of the reads according to the modulation rule. Step 2, obtain the marked read by using MAFFT^26^ to align the observed carrier strand to the carrier strand. Step 3, deduce the consensus sequence for each cluster of marked reads using a simple voting strategy. In the voting process, bases that are marked as insertion, deletion, or substitution errors should not be considered. Finally, the consensus sequences can be demodulated into the binary data by reversing the encoding rule.

## Data availability

All data are available in the main text or the supplementary materials.

## Code availability

Code can be downloaded from https://github.com/BertZan/Modulation-based-DNA-storage

## Acknowledgements

This work was supported by the National Natural Science Foundation of China (grant nos.62072128 and 62002079).

## Author contributions

W.B.L. and P.X. supervised the research. W.B.L. and X.Z.Z. conceived the concept. W.B.L. managed coauthor contributions to the paper. X.Z.Z. wrote the Python codes, performed the simulations and analyzed the data. R.Z.X. polished the paper. R.Z.X. and X.Y.Y. discussed on the data. All authors contributed to the writing of the paper.

## Competing interests

The authors declare no competing interests.

## Competing interests

The authors declare no competing interests.

## Notes

### Competing Interest Statement

The authors have declared no competing interest.

### Summary of Updates

update more comparision results.

## References

1. Ceze, L., Nivala, J. & Strauss, K. Molecular digital data storage using DNA. Nature Reviews Genetics 20, 456–466 (2019).

2. Meiser, L.C. et al. Synthetic DNA applications in information technology. Nature Communications 13, 352 (2022).

3. Takahashi, C.N., Nguyen, B.H., Strauss, K. & Ceze, L. Demonstration of End-to-End Automation of DNA Data Storage. Scientific Reports 9, 4998 (2019).

4. Newman, S. et al. High density DNA data storage library via dehydration with digital microfluidic retrieval. Nature Communications 10, 1706 (2019).

5. Antkowiak, P.L. et al. Integrating DNA Encapsulates and Digital Microfluidics for Automated Data Storage in DNA. Small 18 (2022).

6. Srinivasavaradhan, S.R., Gopi, S., Pfister, H. & Yekhanin, S. Trellis BMA: coded trace reconstruction on IDS channels for DNA storage. (2021).

7. Cretu Stancu, M. et al. Mapping and phasing of structural variation in patient genomes using nanopore sequencing. Nature Communications 8, 1326 (2017).

8. Antkowiak, P.L. et al. Low cost DNA data storage using photolithographic synthesis and advanced information reconstruction and error correction. Nature Communications 11, 5345 (2020).

9. Organick, L. et al. Random access in large-scale DNA data storage. Nat Biotechnol 36, 242–248 (2018).

10. Goldman, N. et al. Towards practical, high-capacity, low-maintenance information storage in synthesized DNA. Nature 494, 77–80 (2013).

11. Meiser, L.C. et al. Reading and writing digital data in DNA. Nat Protoc 15, 86–101 (2019).

12. Davey, M.C. & Mackay, D.J.C. in 2000 IEEE International Symposium on Information Theory (Cat. No.00CH37060) 477 (2000).

13. Lenz, A., Maarouf, I., Welter, L., Wachter-Zeh, A. & Amat, A. Concatenated Codes for Recovery From Multiple Reads of DNA Sequences. (2020).

14. Yazdi, S.M.H.T., Gabrys, R. & Milenkovic, O. Portable and Error-Free DNA-Based Data Storage. Scientific Reports 7(2017).

15. Press, W.H., Hawkins, J.A., Jones, S.K., Schaub, J.M. & Finkelstein, I.J. HEDGES error-correcting code for DNA storage corrects indels and allows sequence constraints. P Natl Acad Sci USA 117, 18489–18496 (2020).

16. Song, L., Geng, F., Gong, Z., Li, B. & Yuan, Y. Super-robust data storage in DNA by de Bruijn graph-based decoding. bioRxiv, 2020.2012.2020.423642 (2020).

17. Zan, X. et al. A Hierarchical Error Correction Strategy for Text DNA Storage. Interdisciplinary Sciences: Computational Life Sciences 14, 141–150 (2022).

18. Xiangzhen ZAN et al. An Efficient Bueket-allocation Decoding Method Based on Forward Error Correction Codes for Deoxyribo Nucleicecid Storage. Journal of Electronics and Information Technology 44, 1–7 (2021).

19. Xue, T.B. & Lau, F.C.M. Construction of GC-Balanced DNA With Deletion/Insertion/Mutation Error Correction for DNA Storage System. Ieee Access 8, 140972–140980 (2020).

20. Levenshtein, V.I. Binary Codes Capable of Correcting Deletions, Insertions, and Reversals. Soviet Physics Doklady 10, 707–710 (1965).

21. Church, G.M., Gao, Y. & Kosuri, S. Next-Generation Digital Information Storage in DNA. Science 337, 1628 (2012).

22. Erlich, Y. & Zielinski, D. DNA Fountain enables a robust and efficient storage architecture. Science 355, págs. 950–954 (2017).

23. Zhan, Y.J. & Huang, F.C. Generalized Spatial Modulation With Multi-Index Modulation. Ieee Commun Lett 24, 585–588 (2020).

24. Moore, B.C. & Sek, A. Effects of carrier frequency, modulation rate, and modulation waveform on the detection of modulation and the discrimination of modulation type (amplitude modulation versus frequency modulation). The Journal of the Acoustical Society of America 97, 2468–2478 (1995).

25. Agrawal, G.P. Modulation instability induced by cross-phase modulation. Physical review letters 59, 880–883 (1987).

26. Katoh, K., Misawa, K., Kuma, K.i. & Miyata, T. MAFFT: a novel method for rapid multiple sequence alignment based on fast Fourier transform. Nucleic Acids Res 30, 3059–3066 (2002).

27. Sievers, F. et al. Fast, scalable generation of high-quality protein multiple sequence alignments using Clustal Omega. Mol Syst Biol 7(2011).

28. Notredame, C., Higgins, D.G. & Heringa, J. T-Coffee: A novel method for fast and accurate multiple sequence alignment. Journal of Molecular Biology 302, 205–217 (2000).

29. Edgar, R.C. MUSCLE: multiple sequence alignment with high accuracy and high throughput. Nucleic Acids Res 32, 1792–1797 (2004).

30. Needleman, S. Needleman-Wunsch algorithm for sequence similarity searches. J Mol Biol 48, 443–453 (1970).

31. Wang, K. et al. MFEprimer-3.0: quality control for PCR primers. Nucleic Acids Res 47, W610–W613 (2019).

32. van Dijk, E.L., Jaszczyszyn, Y., Naquin, D. & Thermes, C. The Third Revolution in Sequencing Technology. Trends Genet 34, 666–681 (2018).

33. Jain, M. et al. MinION Analysis and Reference Consortium: Phase 2 data release and analysis of R9.0 chemistry. F1000Res 6, 760–760 (2017).

34. Rashtchian, et al. Clustering Billions of Reads for DNA Data Storage. Advances in Neural Information Processing Systems 30 (2017).

